# An endolysin-expressing probiotic attenuates *Clostridioides difficile* infection in germ-free and dysbiotic microbiota-associated mice

**DOI:** 10.64898/2026.10.07.757339

**Authors:** Amal Cheemadan, Phurt Harnvoravongchai, Sunil More, Purna C. Kashyap, Joy Scaria

## Abstract

Disruption of the intestinal microbiota increases susceptibility to *Clostridioides difficile* infection (CDI), creating an opportunity for prophylactic interventions before symptomatic disease develops. We investigated whether *Escherichia coli* Nissle 1917 (EcN) engineered to express the bacteriophage-derived endolysin CD27L could enhance pathogen inhibition and attenuate CDI. Using a plasmid-cured EcN chassis and pMUT2-derived vectors, we generated strains expressing CD27L under constitutive or anaerobically regulated promoter control. The constitutive strain, BioEcN1, reduced viable *C. difficile* recovery relative to parental EcN and was selected for animal studies. BioEcN1 remained recoverable from monoassociated mice without *in vivo* antibiotic selection, with similar recovery on selective and nonselective media supporting construct retention. Prophylactic administration reduced clinical disease, body-weight loss, and intestinal histopathological injury in germ-free mice. Observed survival was 100% in BioEcN1-treated challenged mice versus 50% in untreated challenged mice during the monitored period. Protection was accompanied by lower fecal and cecal pathogen burdens. In mice harboring a human-derived dysbiotic microbiota, prophylactic BioEcN1 also reduced clinical and stool-consistency scores, with lower early fecal pathogen recovery and less colon shortening. These findings establish preclinical proof of concept for prophylactic administration of an endolysin-expressing probiotic to attenuate CDI.

## Introduction

*Clostridioides difficile* is a spore-forming, toxin-producing intestinal pathogen that causes disease ranging from diarrhea to life-threatening colitis^1–4^. *C. difficile* infection (CDI) has a substantial healthcare burden, with an estimated 462,100 cases in the United States in 2017^5^. Antibiotic exposure is a principal risk factor for CDI because it disrupts the intestinal microbiota and weakens microbiota-mediated colonization resistance^6–8^. The resulting changes in community composition and metabolism create conditions that support pathogen establishment and proliferation^9^, Older age^10,11^, immunocompromising conditions^12^, and recent hospitalization further increase risk by weakening host defenses and providing opportunities for pathogen exposure^13^. These identifiable risk settings provide a rationale for interventions that can be used before symptomatic infection develops. In this context, a prophylactic strategy that limits pathogen expansion while minimizing further disruption of the resident microbiota could help protect susceptible individuals during this interval.

Microbiota-based therapies have demonstrated that modifying the intestinal community can reduce CDI recurrence after antibacterial treatment^14,15^. Their efficacy supports the value of microbial interventions and motivates investigation of strategies that act before an initial infection. One approach is to introduce a defined probiotic engineered to provide localized antimicrobial activity against *C. difficile*^16^. Persistence without antibiotic selection could help maintain this activity during the susceptible period^17^. Combining intestinal persistence with a pathogen-directed antimicrobial function therefore offers a rationale for developing engineered probiotics for CDI prophylaxis^18^.

*Escherichia coli Nissle 1917* (EcN) provides an established probiotic chassis for developing such an intervention^19^. Its history of probiotic use extends over a century, with clinical investigation supporting its value in specific gastrointestinal conditions^20,21^. For example, in a randomized trial involving 327 patients with ulcerative colitis, EcN showed efficacy comparable to mesalazine for maintaining remission over 12 months, with similar tolerability^22^. Beyond its clinical history, EcN benefits from extensive knowledge of *E. coli* biology and genetic tools that enable the introduction of defined therapeutic functions^23,24^. Engineered EcN has delivered recombinant proteins within the mouse intestine, supporting its use as a local delivery organism^25^. Native-plasmid-derived expression systems have further enabled intestinal recombinant protein production without antibiotic selection for plasmid maintenance^17^. These features make EcN a suitable platform for testing whether an antimicrobial payload can enhance inhibition of *C. difficile* and provide prophylactic protection. Engineered probiotics have already been used to attenuate experimental CDI by modifying intestinal bile salt metabolism, illustrating how defined genetic functions can influence pathogen susceptibility^16^. Equipping EcN with an antimicrobial payload offers another approach: augmenting the parental strain with activity directed against *C. difficile*. Intestinal persistence and functional expression are central requirements for evaluating this strategy.

Bacteriophages provide a source of antibacterial functions that can be incorporated into a probiotic chassis^26,27^. Successful rescue treatment of multidrug-resistant infections has demonstrated the therapeutic potential of intact phages^28,29^, although personalized treatment can require identifying suitable phages and preparing formulations active against the infecting isolate^30^. Phage-derived endolysins offer a way to use a defined antibacterial component without administering the intact virus^31^. These enzymes degrade bacterial peptidoglycan during phage release, and selected endolysins can also lyse susceptible bacteria when applied externally^32^. CD27L, derived from the *C. difficile* bacteriophage ΦCD27, exhibited activity against 30 diverse *C. difficile* strains while sparing the commensal species examined in the same study^27^. These findings support its evaluation as a pathogen-directed antimicrobial payload. Producing CD27L in an intestinal probiotic could make antimicrobial activity available before pathogen exposure and during early colonization. The intended benefit is to limit pathogen expansion during impaired colonization resistance, before toxin-mediated injury and symptomatic disease develop. This preventive strategy depends on the engineered organism persisting and producing active endolysin within the intestinal environment. Whether CD27L expression enhances EcN-mediated inhibition and provides protection in the presence of a dysbiotic microbial community remains unresolved. We therefore evaluated CD27L-engineered EcN as a prophylactic intervention against *C. difficile*.

Here, we generated plasmid-cured EcN strains carrying pMUT2-derived CD27L expression constructs under constitutive or anaerobically regulated promoter control. We compared their activity against multiple *C. difficile* strains and selected BioEcN1 for animal experiments based on its enhanced coculture inhibition relative to parental EcN. We then evaluated bacterial recovery without *in vivo* antibiotic selection and prophylactic protection in germ-free mice and mice harboring a human-derived dysbiotic microbiota. BioEcN1 administration before challenge reduced clinical disease in both models and limited intestinal injury in germ-free mice. These findings support a defined engineered-probiotic approach to attenuating CDI during a period of microbiota-associated susceptibility.

## 2. Materials and methods

### 2.1. Study design

This study evaluated whether plasmid-cured *Escherichia coli* Nissle 1917 (EcN) could be engineered to express the bacteriophage-derived endolysin CD27L and inhibit *C. difficile*. We first constructed EcN strains expressing CD27L from a constitutive J23101 promoter (BioEcN1) or an anaerobically regulated FNRS promoter(BioEcN2). These strains were evaluated under anaerobic coculture conditions to assess their ability to inhibit *C. difficile* growth.

BioEcN1 was advanced to animal studies based on its more consistent inhibitory activity across the tested *C. difficile* strains. We then assessed BioEcN1 colonization and recombinant plasmid retention in germ-free mice before testing prophylactic administration in germ-free and dysbiotic microbiota-associated models of *C. difficile* infection (CDI). Wild-type EcN was included in the in vitro studies but was not included as a separate animal group because it did not significantly inhibit *C. difficile* under the relevant *in vitro* conditions. Omitting this group reduced animal use in accordance with the reduction principle of the 3Rs.

### 2.2. Bacterial strains, plasmids, and culture conditions

EcN was used as the probiotic chassis. *E. coli* DH5alpha was used for plasmid propagation, and *E. coli* BL21(DE3) was used for pET28a-based protein-expression studies. The *C. difficile* strains used in the inhibition assays were R20291, strain 630 (ATCC BAA-1382), and VPI 10463 (ATCC 43255). *E. coli* strains were routinely cultured in LB Miller broth or on LB agar at 37^₀^C unless otherwise indicated. Ampicillin (100 µg/mL), kanamycin (50 µg /mL), or the antibiotic appropriate for each plasmid was added when selection was required. *C. difficile* cultures were maintained at 37^₀^C under anaerobic conditions in pre-reduced modified brain heart infusion (mBHI) medium^33^. All strains, plasmids, resistance markers, and relevant genotypes are listed in Supplementary Table S1.

### 2.3. Removal of native EcN plasmids and preparation of electrocompetent cells

The native EcN plasmids pMUT1 and pMUT2 were removed using the CRISPR-Cas9 plasmid-curing system described by Kan et al.^17^. EcN was transformed with pFREE to remove pMUT1 and with pCryptDel4.8 to remove pMUT2. Following each transformation, cells were cultured overnight in LB containing 50 micrograms/mL kanamycin. Cultures were diluted 1:1,000 into fresh LB containing 50 micrograms/mL kanamycin, 0.2% rhamnose, and 0.43 micromolar anhydrotetracycline and incubated at 37^₀^C for 8 hours. Cultures were streaked on LB agar without antibiotics and incubated overnight. Colonies lacking the native plasmids were identified by colony PCR using primers muta5, muta6, muta7, and muta8. The resulting plasmid-cured strain is referred to as deltaEcN.

Electrocompetent deltaEcN cells were prepared from overnight cultures diluted into 1.4 mL LB and incubated at 37^₀^C with shaking at 900 rpm. After approximately 2 hours, when the OD600 reached approximately 0.6, cells were collected at 9,000 rpm at 24^₀^C. The pellet was washed twice with 1 mL sterile deionized water at room temperature and resuspended in approximately 30 microliters water. Plasmid DNA or assembly product (300 ng) was combined with the cells and electroporated in a 1-mm-gap cuvette at 1,250 V. Cells were recovered in 1 mL fresh medium at 37^₀^C for 1 hour and plated on LB agar containing the appropriate antibiotic.

### 2.4. Construction of CD27L expression plasmids

The coding sequence for the phiCD27 endolysin CD27L was selected based on its previously reported activity against *C. difficile* ^27^. The sequence was codon optimized for expression in *E. coli*. An OmpA signal peptide (MKKTAIAIAVALAGFATVAQA) was placed at the N terminus, and a C-terminal polyhistidine tag was included for protein detection. The complete expression cassette contained a ribosome-binding site, the OmpA-CD27L coding sequence, and a transcriptional terminator. Expression was controlled by either the constitutive J23101 promoter or the anaerobically regulated FNRS promoter.

The synthetic CD27L cassette was initially obtained in pUC19. To generate the EcN expression plasmids, the J23101-CD27L or FNRS-CD27L insert and the pMUT2-derived backbone were amplified with Q5 High-Fidelity DNA Polymerase (New England Biolabs). Primers were designed in SnapGene to introduce the overlaps required for assembly. Amplicons were resolved on 1% agarose gels, and bands of the expected size were purified using the GeneJET Gel Extraction Kit (Thermo Fisher Scientific; catalog no. K0691). Purified inserts and backbone were combined at molar ratios and assembled using NEBuilder HiFi DNA Assembly Master Mix. Reactions were incubated at 50^₀^C for 15 minutes.

Assembly products were transformed into chemically competent *E. coli* DH5alpha, recovered in SOC medium, and plated on LB agar containing 100 micrograms/mL ampicillin. Candidate colonies were screened by colony PCR with primers spanning the CD27L cassette and pMUT2-backbone junctions. Colony-PCR products were resolved on 1% TAE agarose gels at 120 V for 35 minutes and imaged with an AAB Advanced American Biotechnology UV transilluminator or ChemiDoc™ MP Imaging System (Bio-Rad Laboratories, Inc., Hercules, CA, USA). Confirmed plasmids were introduced into deltaEcN to generate BioEcN1 and BioEcN2.

For independent validation of CD27L protein production, the 873-bp OmpA-CD27L fragment was amplified from the pUC19 construct with primers introducing NcoI and SacI sites. The amplicon and pET28a vector were digested with NcoI and SacI, gel purified, and ligated with T4 DNA Ligase (New England Biolabs; catalog no. M0202). The ligation product was transformed into chemically competent DH5alpha, and transformants were selected on LB agar containing 50 micrograms/mL kanamycin. Candidate colonies were confirmed by colony PCR before transfer of the plasmid into BL21(DE3). Separate reporter constructs fused superfolder green fluorescent protein (sfGFP) to CD27L through a GGGSGGG linker.

### 2.5. Confirmation of CD27L expression

#### SDS-PAGE analysis

BL21(DE3) carrying pET28a-CD27L was cultured to an OD600 of approximately 0.5 and induced with 0.1 mM isopropyl beta-D-1-thiogalactopyranoside. Cultures were incubated for 10 hours at 20^₀^C with shaking at 150 rpm. Cells were collected at 13,000 rpm for 5 minutes, resuspended in phosphate-buffered saline (PBS), and lysed by five freeze-thaw cycles. Lysates were clarified by centrifugation, and protein concentrations were determined by Bradford assay.

Clarified lysates were combined with 4x Laemmli sample buffer at a 3:1 sample-to-buffer ratio and heated at 95^₀^C for 5 minutes. Proteins were separated using a 4% stacking gel and 12% resolving gel in Tris-glycine-SDS buffer. Electrophoresis was performed at 80 V through the stacking gel and 120 V through the resolving gel. A prestained 10- to 250-kDa protein ladder (New England Biolabs; P7719S) was used as a molecular-mass marker. Gels were stained with 0.2% Coomassie Brilliant Blue R-250 in 50% methanol and 37.5% acetic acid and destained in 10% methanol and 0.75% acetic acid. CD27L expression was evaluated from the presence of a band at the expected molecular mass of approximately 32 kDa.

#### Fluorescence microscopy

Reporter plasmids encoding CD27L-sfGFP were introduced into deltaEcN. Single colonies carrying the J23101- or FNRS-controlled reporter constructs were cultured under the conditions used to evaluate the corresponding promoter. Cells were imaged using a Motic BA410E fluorescence microscope. Matched bright-field and GFP images were collected for each construct and its nonfluorescent control.

### 2.6. *In vitro* inhibition assays

Cultures of *C. difficile* R20291, strain 630, or VPI 10463 and the indicated EcN strain were standardized to an optical density of 0.5 measured at 630nm. Equal volumes of the *C. difficile* and EcN suspensions (500µL) were combined in pre-reduced mBHI broth. Untreated *C. difficile*, wild-type EcN, BioEcN1, and BioEcN2 conditions were included for each pathogen strain. Cocultures were incubated anaerobically at 37^₀^C for 24 hours. Following incubation, cultures were serially diluted and plated on *C. difficile*-selective agar consisting of mBHI agar supplemented with D-cycloserine at 0.5 g/L and cefoxitin at 0.008 g/L. Plates were incubated anaerobically at 37^₀^C for 24 hours before colonies were enumerated. Counts were reported as CFU/mL. Each assay included five replicate cultures per condition and was performed in two independent experiments.

### 2.7. Animals, husbandry, and ethical oversight

The germ-free C57BL/6 breeding colony was established using mice obtained from Taconic Biosciences (New York, USA). Experimental mice were bred and maintained in the Oklahoma State University Germ-Free Mouse Core Facility, operated by our laboratory, and enrolled in the study at five weeks of age. Mice were housed in an Allentown SPP system under a 12-hour light-dark cycle and received sterilized standard chow and autoclaved water *ad libitum*. Bedding, food, water, and experimental materials were sterilized before introduction into the gnotobiotic housing system. Animals were assigned to experimental groups before treatment. Histopathological evaluation was performed by an observer blinded to treatment group. All animal procedures were approved by the Oklahoma State University Institutional Animal Care and Use Committee under protocol IACUC-25-30-STW and followed institutional and federal guidelines for humane animal use.

### 2.8. BioEcN1 colonization and plasmid retention in germ-free mice

BioEcN1 was grown to late logarithmic phase (OD600 approximately 1.0), collected by centrifugation, and resuspended in sterile PBS. Germ-free mice received approximately 1 x 10^8 CFU BioEcN1 by oral gavage on day 0. Fecal pellets were collected longitudinally for 10 days, and body weight was recorded daily. Fresh fecal pellets were weighed and homogenized in sterile PBS. Homogenates were serially diluted and plated on LB agar without ampicillin to estimate total recoverable EcN and on LB agar containing 100 micrograms/mL ampicillin to estimate BioEcN1 retaining the recombinant plasmid. Plates were incubated at 37^₀^C before CFU enumeration.. Ampicillin was used only for analytical recovery; mice did not receive antibiotic selection.

### 2.9. Prophylactic BioEcN1 administration in germ-free mice

Germ-free mice were assigned to four groups: uninfected germ-free control, BioEcN1 only, CDI, and BioEcN1 plus CDI. Each group contained 5-6 mice. Mice assigned to BioEcN1 received approximately 1 x 10^8 CFU by oral gavage on days 0 and 6. Control animals received an equivalent volume of sterile PBS. On day 7, mice in the infection groups were orally challenged with approximately 1 x 10^6 CFU *C. difficile* R20291 in sterile PBS; uninfected animals received PBS. Body weight, survival, clinical condition, and stool consistency were assessed daily after challenge. Fecal pellets were collected at the designated time points, including 24 hours after challenge, for enumeration of *C. difficile*. Mice were euthanized on day 12 or when predefined humane-endpoint criteria were reached. At necropsy, cecal contents, cecum, and colon were collected. Cecal contents were used for *C. difficile* enumeration; cecum weight and colon length were recorded; and colon tissue was allocated for histopathology, periodic acid-Schiff staining, and RNA analysis.

### 2.10. Establishment of human-derived dysbiotic microbiota-associated mice

Dysbiotic microbiota-associated donor mice were maintained in the Kashyap laboratory at Mayo Clinic as described previously by Battaglioli et al.^34^. These donor mice harbored a human-derived, diarrhea-associated dysbiotic fecal microbiota characterized in that study. Frozen fecal pellets collected from the donor mice were shipped to Oklahoma State University and used to colonize germ-free C57BL/6 recipient mice by oral gavage. Fecal material was transferred to an anaerobic chamber and combined with an equal volume of sterile, pre-reduced PBS. PBS was autoclaved, passed through a 0.22-µm filter, and equilibrated under anaerobic conditions before use. The suspension was homogenized, and large particles were removed. The nonsettling fraction was diluted 1:2.5 in pre-reduced PBS and maintained with minimal headspace until administration. Recipient mice received 200 microliters of the suspension by oral gavage. Recipient mice were maintained for 21 days to allow establishment of the transferred community before BioEcN1 administration. Because susceptibility was conferred by the transferred dysbiotic microbiota, no antibiotic pretreatment was administered before *C. difficile* challenge.

### 2.11. BioEcN1 administration to dysbiotic microbiota-associated mice

After establishment of the dysbiotic microbiota, mice received BioEcN1 by oral gavage at approximately 1 x 10^8 CFU per mouse and were subsequently provided drinking water containing 5 x 10^8 CFU/mL BioEcN1. Control mice received untreated water and PBS gavage. Body weight was monitored throughout the colonization period. Fecal pellets were collected at the designated time points and plated on LB agar with and without 100µg /mL ampicillin.

### 2.12. Prophylactic BioEcN1 administration in dysbiotic microbiota-associated mice

Dysbiotic microbiota-associated mice were assigned to the following groups: Control, BioEcN1 pre-colonized, CDI control, and BioEcN1-treated CDI. BioEcN1 was administered prophylactically as described above. Control mice received PBS and untreated drinking water. Mice were challenged by oral gavage with approximately 1 x 10^7 CFU *C. difficile* strain 630 (ATCC BAA-1382) without preceding antibiotic treatment. Clinical condition, stool consistency, and body weight were monitored after challenge. Fecal pellets collected on days 1 and 2 after infection were used to enumerate *C. difficile*. Mice were euthanized on day 7 after challenge. Cecum weight, colon length, and gross intestinal morphology were recorded at necropsy.

### 2.13. Clinical assessment and necropsy

Mice were weighed and examined at least once daily. Clinical disease was assessed using a composite score that included (activity, posture, coat condition, diarrhea, and eye/nose appearance. Each parameter was scored from 0 to 3, with 0 representing normal appearance or behavior and increasing scores indicating greater disease severity). Stool consistency was evaluated using rubric developed by Battaglioli et al. ^34^. Animals meeting predefined humane-endpoint criteria were euthanized and recorded for survival analysis. At scheduled necropsy, the gastrointestinal tract was removed, photographed, and processed immediately. Cecum weight was measured with retained contents, and colon length was measured using a consistent anatomical definition.

### 2.14. Enumeration of EcN and C. difficile

Fecal pellets and cecal contents were weighed, suspended in sterile PBS, and homogenized. Serial dilutions were prepared in PBS. EcN-associated CFU were enumerated on LB agar with or without ampicillin as described above. *C. difficile* was enumerated on pre-reduced selective mBHI agar containing D-cycloserine and cefoxitin. Selective plates were incubated anaerobically at 37^₀^C for 24 hours before colonies were counted. Results were expressed as CFU/mL

### 2.15. Histopathology and periodic acid-Schiff staining

Colon tissues were fixed in neutral buffered formalin, embedded in paraffin, and sectioned at 10 µm. Sections were stained with hematoxylin and eosin and were blindly scored for mucosal ulceration, crypt destruction, lamina propria inflammatory infiltrates, edema or fibrin deposition, epithelial necrosis, perivascular inflammation, and submucosal inflammation. Each feature was scored from 1 to 4 according to the percentage of tissue involved: 1, minimal (0 to 10%); 2, mild (10 to 40%); 3, moderate (40 to 70%); and 4, severe (greater than 70%). Individual feature scores were summed to obtain a cumulative injury score. Additional sections were stained with periodic acid-Schiff reagent to visualize goblet cells and mucin-associated structures.. Images were acquired using an Olympus BX53 microscope equipped with a DP27 digital camera (Olympus). H&E-stained sections were imaged at 200× total magnification using a 20× objective and 10× ocular lens, whereas PAS-stained sections were imaged at 100× total magnification using a 10× objective and 10× ocular lens. Images were captured using cellSens software and processed in Adobe Photoshop for white-balance adjustment and annotation. Representative images were selected without knowledge of treatment group.

### 2.16. Statistical analysis

Statistical analyses were performed using GraphPad Prism 10 (GraphPad Software, Boston, MA, USA). Each *in vitro* assay included five replicate cultures per condition and was performed in two independent experiments. Two-group comparisons were analyzed using an unpaired two-tailed Student’s ***t***-test, whereas comparisons involving more than two groups were analyzed using one-way ANOVA. Measurements evaluated across multiple groups and time points were analyzed using two-way ANOVA. Survival was analyzed using Kaplan–Meier survival curves, and differences between groups were assessed using the log-rank. Histopathology scores were analyzed using one-way ANOVA. Data are presented as mean ± SEM. Statistical significance was defined as *P* < 0.05, with significance indicated as *P < 0.05, **P < 0.01, ***P < 0.001, and ****P < 0.0001; ns, not significant

## 3. Results

### Engineering EcN for CD27L expression under constitutive and anaerobically regulated promoters

To develop a probiotic with activity against *C. difficile*, we engineered *E. coli* Nissle 1917 (EcN) to express the bacteriophage-derived endolysin CD27L. We removed the native plasmids pMUT1 and pMUT2 using the previously published curing procedure of Kan et al. (2021), generating a plasmid-cured recipient designated ΔEcN. We then introduced pMUT2-derived CD27L expression constructs into this recipient to generate BioEcN1 and BioEcN2 (Figure 1a).

**Figure 1.**
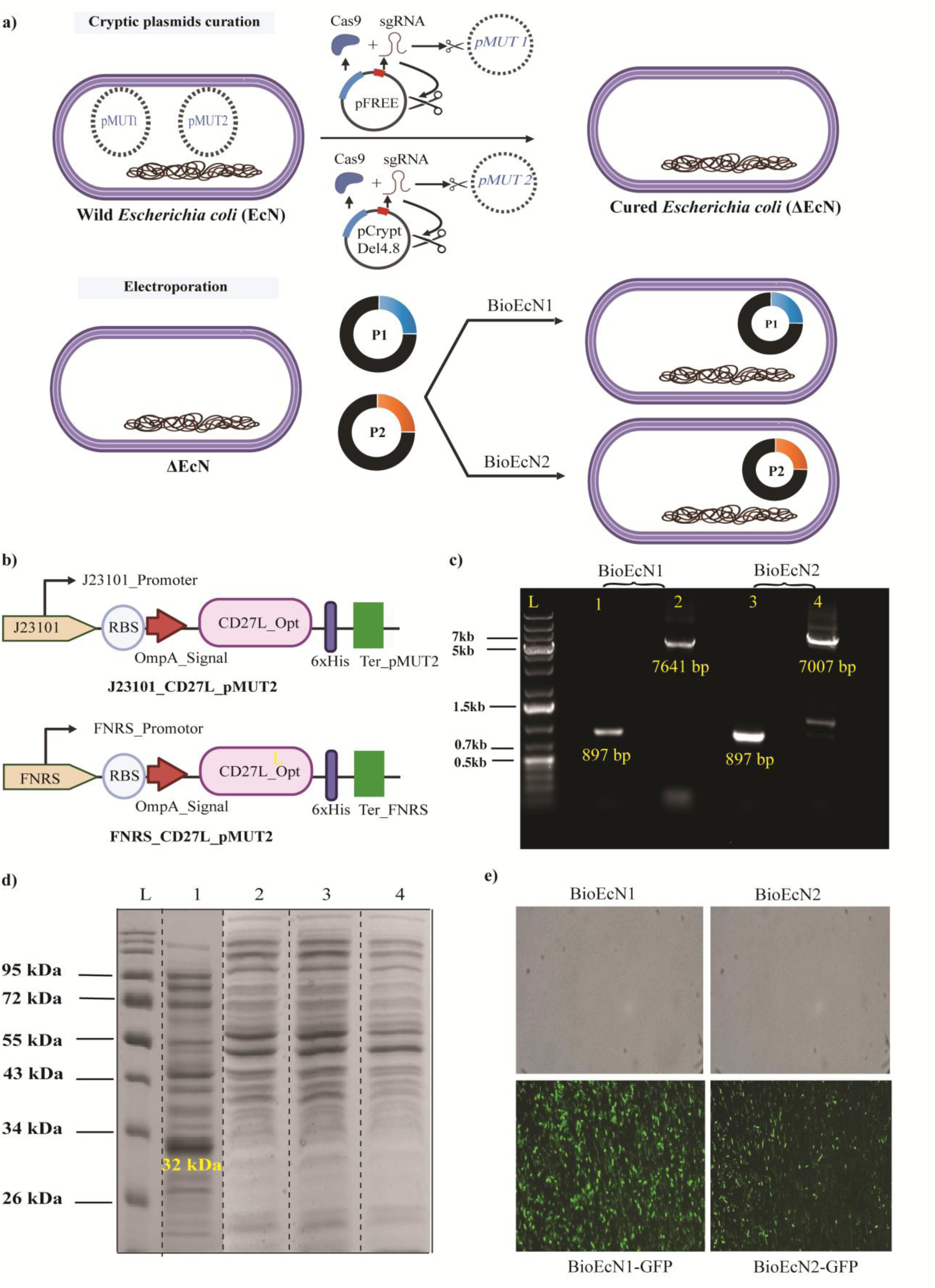
Construction and expression analysis of CD27L-engineered *EcN*. (a) Engineering workflow showing removal of the native plasmids pMUT1 and pMUT2 from *E. coli* Nissle 1917 (EcN), followed by introduction of pMUT2-derived CD27L expression constructs into the plasmid-cured recipient, ΔEcN. Plasmid curing was performed using the previously described method of Kan et al. (2021). (b) Expression cassettes containing the constitutive J23101 or anaerobically regulated FNRS promoter, ribosome-binding site (RBS), OmpA signal peptide, CD27L coding sequence, and C-terminal hexahistidine tag. The corresponding engineered strains were designated BioEcN1 and BioEcN2, respectively. (c) Agarose gel analysis of DNA fragments used in construct generation (d) Coomassie-stained SDS-PAGE analysis of the separate pET28a-CD27L/BL21(DE3) expression system. The arrow indicates a band near the expected molecular mass of approximately 32 kDa. (e) Representative bright-field and fluorescence images of ΔEcN carrying the corresponding CD27L–sfGFP reporter constructs. sfGFP, superfolder green fluorescent protein.

The constructs placed codon-optimized CD27L under the constitutive J23101 promoter or the anaerobically regulated FNRS promoter. Both expression cassettes included a ribosome-binding site, an N-terminal OmpA signal peptide, and a C-terminal hexahistidine tag (Figure 1b). Agarose gel analysis of DNA fragments used in construct generation is shown in Figure 1c. Expression of CD27L was verified and SDS-PAGE revealed a band near 32 kDa, consistent with the expected molecular mass of the recombinant CD27L product (Figure 1d). To examine expression in ΔEcN, we evaluated corresponding CD27L–sfGFP reporter constructs. Fluorescence microscopy detected GFP-associated signals in cells carrying each promoter-controlled construct under the tested conditions (Figure 1e). These observations supported reporter expression in the plasmid-cured EcN chassis.

### BioEcN1 enhances inhibition of multiple *C. difficile* strains *in vitro*

Next, we examined whether CD27L-engineered EcN exhibited enhanced inhibitory activity against *C. difficile*. Viable pathogen recovery was compared following anaerobic coculture with parental EcN, BioEcN1, or BioEcN2. Across all three tested strains, BioEcN1 significantly reduced *C. difficile* recovery relative to both untreated cultures and cultures containing parental EcN (Figure 2a–c). Parental EcN also reduced pathogen recovery relative to untreated cultures in two of the three assays (Figure 2a,c), indicating that BioEcN1 enhanced an existing inhibitory effect under these conditions. BioEcN2 significantly reduced pathogen recovery relative to untreated cultures in all three assays, but its activity did not differ significantly from that of parental EcN (Figure 2a–c). BioEcN1 consistently yielded the lowest mean pathogen counts and showed enhanced inhibition relative to parental EcN across the tested strains. These findings supported its selection for subsequent evaluation of intestinal persistence and prophylactic activity in mice.

**Figure 2.**
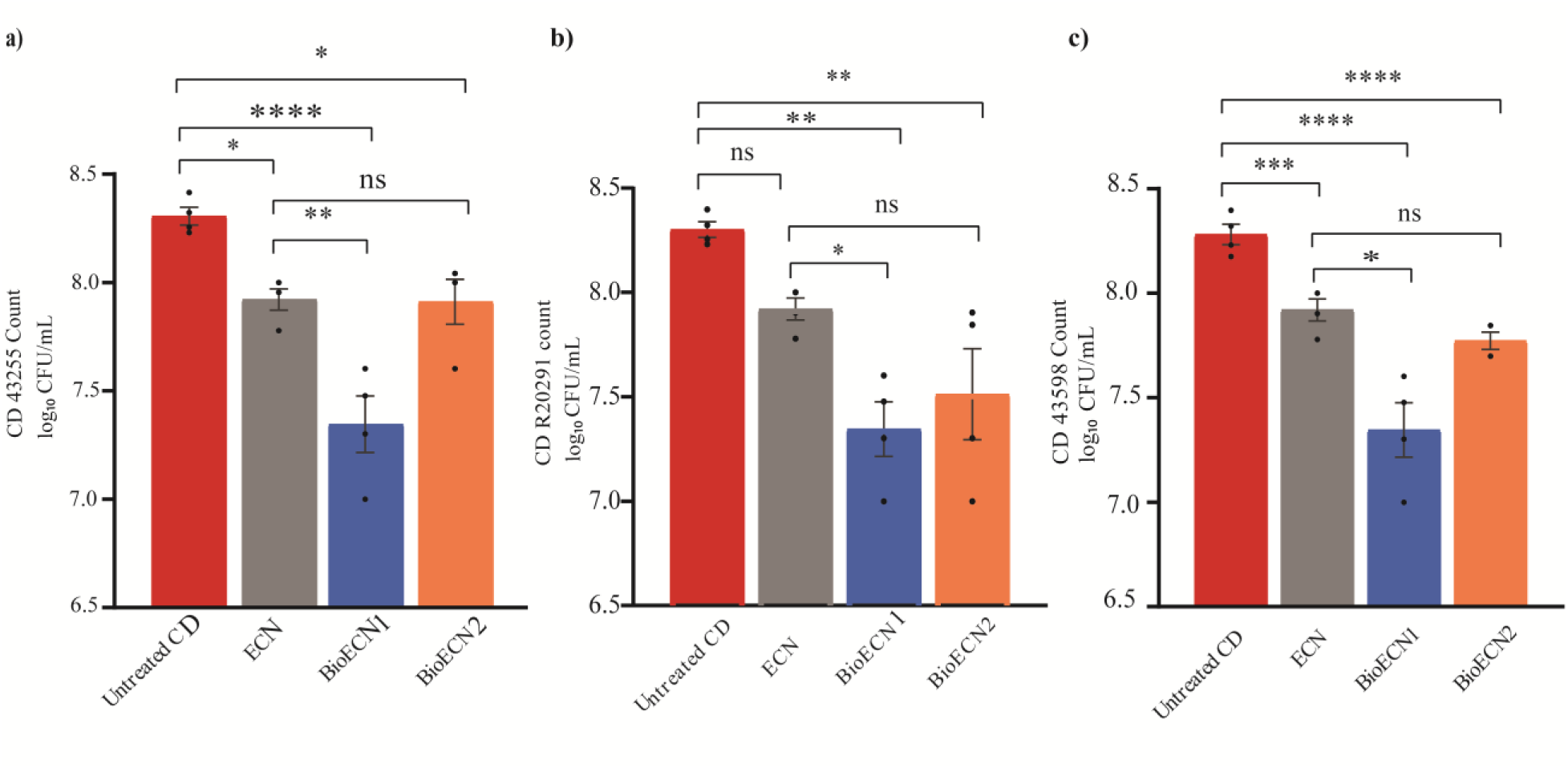
In vitro inhibition of three C. difficile strains by engineered EcN. Viable *C. difficile* recovery following 24-hour anaerobic coculture at 37°C with parental EcN, BioEcN1, or BioEcN2. Cultures containing *C. difficile* alone served as untreated controls. Pathogen and EcN suspensions were standardized and combined in equal volumes before incubation. Panels show strains labeled (a) CD43255, (b) R20291, and (c) CD43598 in the current figure. Pathogen counts were determined by selective plating and expressed as log₁₀ CFU/mL. Each assay included five replicate cultures per condition and was performed in two independent experiments.

### BioEcN1 persists in germ-free mice without *in vivo* antibiotic selection

To assess intestinal persistence and recombinant plasmid retention, we administered BioEcN1 to germ-free mice and monitored fecal bacterial recovery for 10 days(Figure 3a). Fecal suspensions were plated on media with and without ampicillin; mice received no antibiotic selection. BioEcN1 remained recoverable throughout the observation period, with counts generally ranging from approximately 10⁸ to 10⁹ CFU/mL of fecal suspension (Figure 3b). Recovery on ampicillin-containing and nonselective media followed similar trend, supporting retention of the selectable construct during intestinal persistence. During the three-day body-weight assessment, BioEcN1-monoassociated mice maintained mean weights near or above baseline, with a trend similar to that of germ-free controls (Figure 3c). These findings demonstrate stable BioEcN1 recovery without *in vivo* antibiotic selection and no observed decline in mean body weight during the measured period.

**Figure 3.**
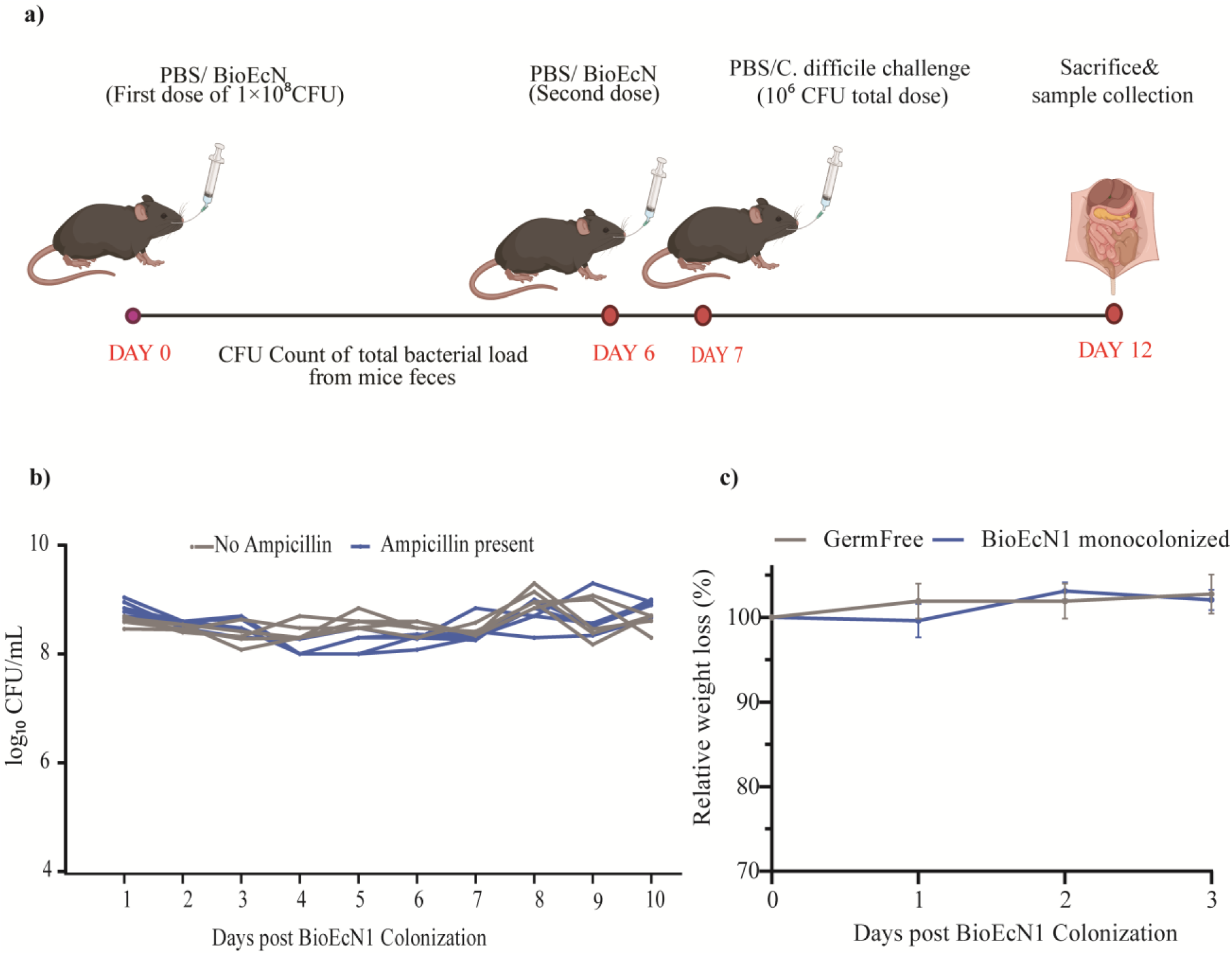
Fecal recovery and body-weight monitoring following BioEcN1 monoassociation. Germ-free C57BL/6 mice received approximately 10⁸ CFU BioEcN1 by oral gavage and were monitored without antibiotic administration. (a) Longitudinal fecal bacterial recovery over 10 days. Fecal suspensions were plated on LB agar without ampicillin and LB agar containing 100 µg/mL ampicillin. Counts are expressed as log₁₀ CFU/mL of fecal suspension. Ampicillin was used only during analytical recovery. (b) Relative body weight of BioEcN1-monoassociated mice and germ-free controls during the three-day assessment, expressed as a percentage of each animal’s baseline weight.

### Prophylactic BioEcN1 attenuates CDI in germ-free mice

In the next step, we evaluated whether BioEcN1 administration before *C. difficile* challenge attenuated disease in germ-free mice. CDI-only mice developed pronounced perianal soiling, whereas mice receiving prophylactic BioEcN1 exhibited a less pronounced stool phenotype (Figure 4a). Clinical scores were significantly lower in BioEcN1-treated challenged mice than in CDI-only mice (Figure 4b). This reduction in clinical disease was accompanied by preservation of body weight. Mean body weight in the CDI-only group declined to approximately 80% of baseline by day 2 after challenge, whereas BioEcN1-treated challenged mice maintained mean weights near baseline through day 3 (Figure 4c). Observed survival declined to 50% in the CDI-only group by approximately 52 hours, while all BioEcN1-treated challenged mice survived throughout the displayed observation period (Figure 4d). Fecal and cecal *C. difficile* recovery was significantly lower in BioEcN1-treated challenged mice than in CDI-only mice (Figure 4e,f). Mean counts differed by approximately 0.3–0.4 log₁₀ CFU/mL of sample suspension, indicating reduced pathogen recovery despite residual colonization. These findings show that prophylactic BioEcN1 attenuated clinical disease in germ-free mice, with preservation of body weight, higher observed survival, and reductions in fecal and cecal pathogen counts.

**Figure 4.**
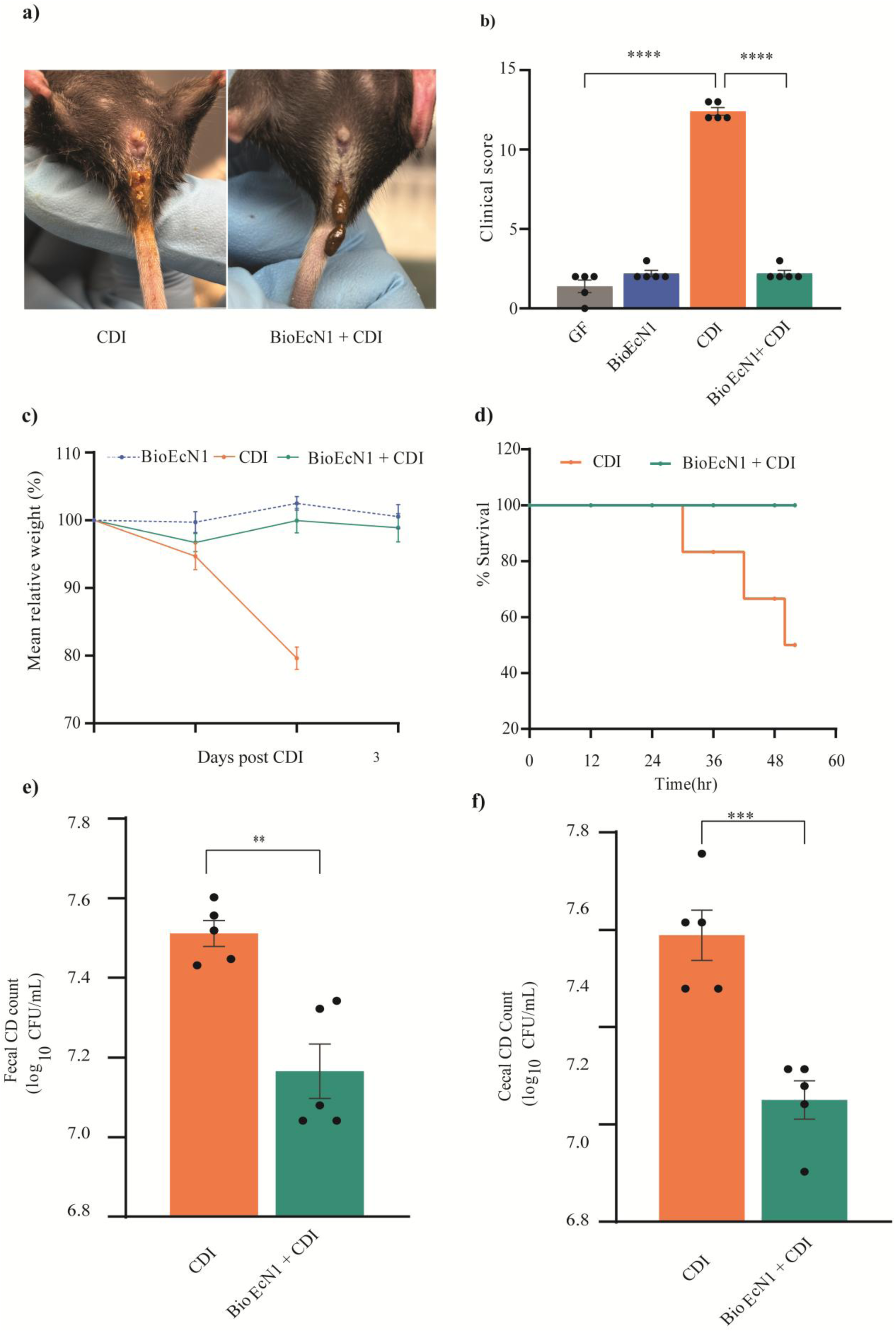
Clinical outcomes and pathogen recovery following prophylactic BioEcN1 administration in germ-free mice. Germ-free C57BL/6 mice received approximately 10⁸ CFU BioEcN1 on days 0 and 6 and were challenged with approximately 10⁶ CFU *C. difficile* R20291 on day 7. Groups comprised uninfected germ-free controls (GF), BioEcN1 alone, *C. difficile* infection alone (CDI), and BioEcN1 followed by challenge (BioEcN1 + CDI). (a) Representative perianal and stool phenotypes in challenged mice. (b) Clinical disease scores. (c) Relative body weight, expressed as a percentage of each animal’s weight immediately before challenge. (d) Kaplan–Meier survival curves over the displayed observation period. (e) Fecal and (f) cecal *C. difficile* counts, expressed as log₁₀ CFU/mL of sample suspension.

### BioEcN1 limits CDI-associated intestinal injury

Intestinal tissue assessment showed that the clinical protection associated with prophylactic BioEcN1 administration extended to reduced tissue injury. Gross examination revealed differences in cecal and colonic appearance between CDI-only mice and mice receiving BioEcN1 before challenge (Figure 5a). Colons were significantly longer in BioEcN1-treated challenged mice than in CDI-only mice, indicating attenuation of infection-associated colon shortening (Figure 5b). Cecum weight was also significantly lower in the BioEcN1-treated challenge group (Figure 5c). Histological assessment supported these findings. Representative H&E-stained sections are shown in Figure 5d. Histopathological injury scores were significantly lower in BioEcN1-treated challenged mice than in CDI-only mice and did not differ significantly from those of germ-free controls (Figure 5e). Together, the anatomical measurements and histopathological scores indicate that prophylactic BioEcN1 administration reduced CDI-associated intestinal injury.

**Figure 5.**
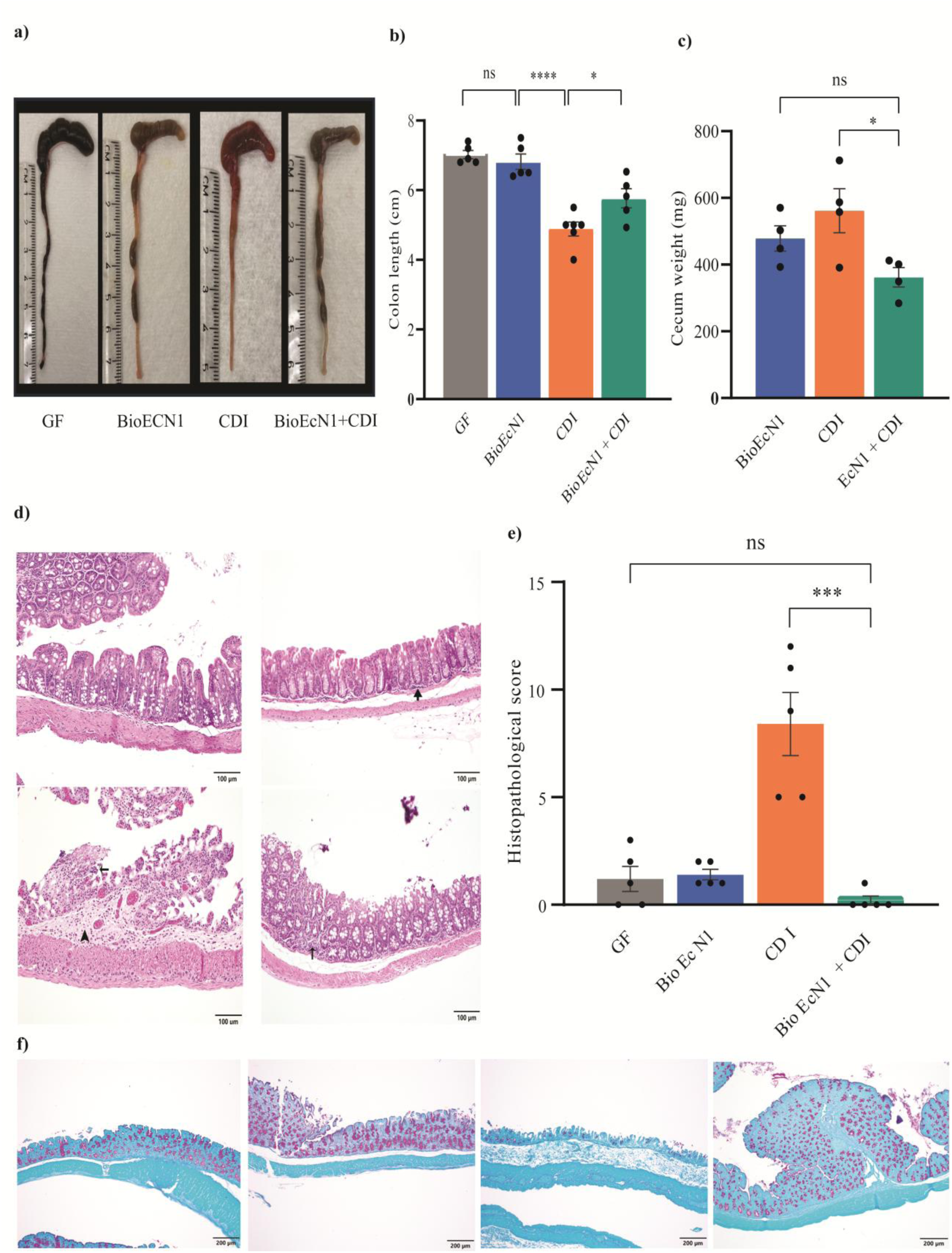
Gross and histological assessment of intestinal injury following BioEcN1 prophylaxis. Intestinal tissues were examined in uninfected germ-free controls (GF), mice receiving BioEcN1 alone, mice challenged with *C. difficile* R20291 (CDI), and mice receiving BioEcN1 before challenge (BioEcN1 + CDI). (a) Representative gross images of ceca and colons. Rulers indicate centimeters. (b) Colon length. (c) Cecum weight [specify whether contents were included]. (d) Representative hematoxylin-and-eosin-stained colon sections. (e) Histopathological injury scores assessed by a pathologist blinded to treatment. (f) Representative colon section. Scale bars, 200 µm. Points in b, c, and e represent individual mice.

### BioEcN1 recovery in mice harboring a human-derived dysbiotic microbiota

To evaluate BioEcN1 recovery in the presence of a complex microbial community, we colonized germ-free recipients with fecal material from donor mice harboring a human-derived dysbiotic microbiota. Following community establishment, mice received BioEcN1 by oral gavage and through drinking-water supplementation. Fecal samples were collected over seven days to assess bacterial recovery (Figure 6a). Bacteria were recovered on both nonselective and ampicillin-containing media at days 1, 3, and 7 after BioEcN1 administration (Figure 6b). Counts on ampicillin-containing plates remained approximately 10⁸–10⁹ CFU/mL of fecal suspension. Recovery under ampicillin selection was consistent with the presence of BioEcN1. Mean body weight remained near or above baseline in both BioEcN1-treated mice and dysbiotic microbiota-associated controls during the seven-day observation period (Figure 6c). These findings support recovery of ampicillin-resistant bacteria during BioEcN1 administration without an accompanying decline in mean body weight.

**Figure 6.**
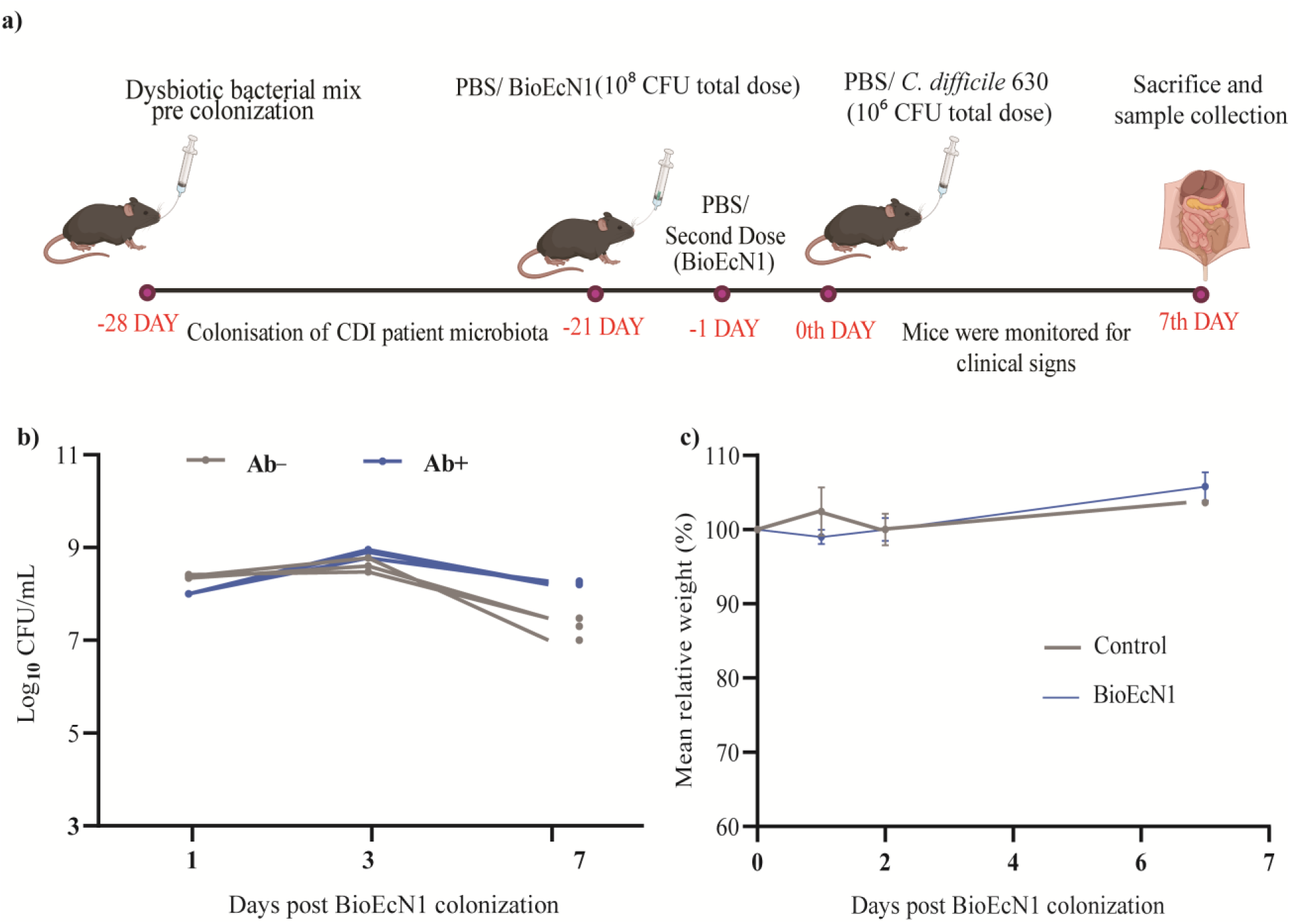
Bacterial recovery and body-weight monitoring during BioEcN1 administration in dysbiotic microbiota-associated mice. (a) Experimental timeline showing transfer of fecal microbiota from dysbiotic microbiota-associated donor mice, community establishment in germ-free C57BL/6 recipients, BioEcN1 administration, and fecal sampling. BioEcN1 was administered by oral gavage and through drinking-water supplementation. (b) Bacterial recovery from fecal suspensions collected on days 1, 3, and 7 after initial BioEcN1 administration. Samples were plated on LB agar without ampicillin or with 100 µg/mL ampicillin, and counts are expressed as log₁₀ CFU/mL of fecal suspension. (c) Relative body weight of BioEcN1-treated mice and dysbiotic microbiota-associated controls, expressed as a percentage of each animal’s baseline weight.

### Prophylactic BioEcN1 attenuates CDI in dysbiotic microbiota-associated mice

Finally, we evaluated whether prophylactic BioEcN1 administration attenuated CDI in mice harboring a human-derived dysbiotic microbiota. Following challenge with *C. difficile* strain 630, BioEcN1-treated mice had significantly lower clinical scores than CDI-only mice (Figure 7a). Stool-consistency scores were also significantly lower, indicating reduced diarrhea severity (Figure 7b). Reduced clinical disease was accompanied by lower early pathogen recovery. Mean fecal *C. difficile* counts were approximately 1.3–1.4 log₁₀ CFU/mL lower in BioEcN1-treated mice than in CDI-only mice on days 1 and 2 after challenge, although the pathogen remained detectable in both groups (Figure 7c). At necropsy, colons were significantly longer in BioEcN1-treated challenged mice than in CDI-only mice (Figure 7d), with representative intestinal images shown in Figure 7e. Cecum weights were also significantly lower in the treated group (Figure 7f), accompanied by differences in gross cecal appearance (Figure 7g). Together, these findings show that prophylactic BioEcN1 administration attenuated clinical disease and intestinal abnormalities in mice harboring a human-derived dysbiotic microbial community.

**Figure 7.**
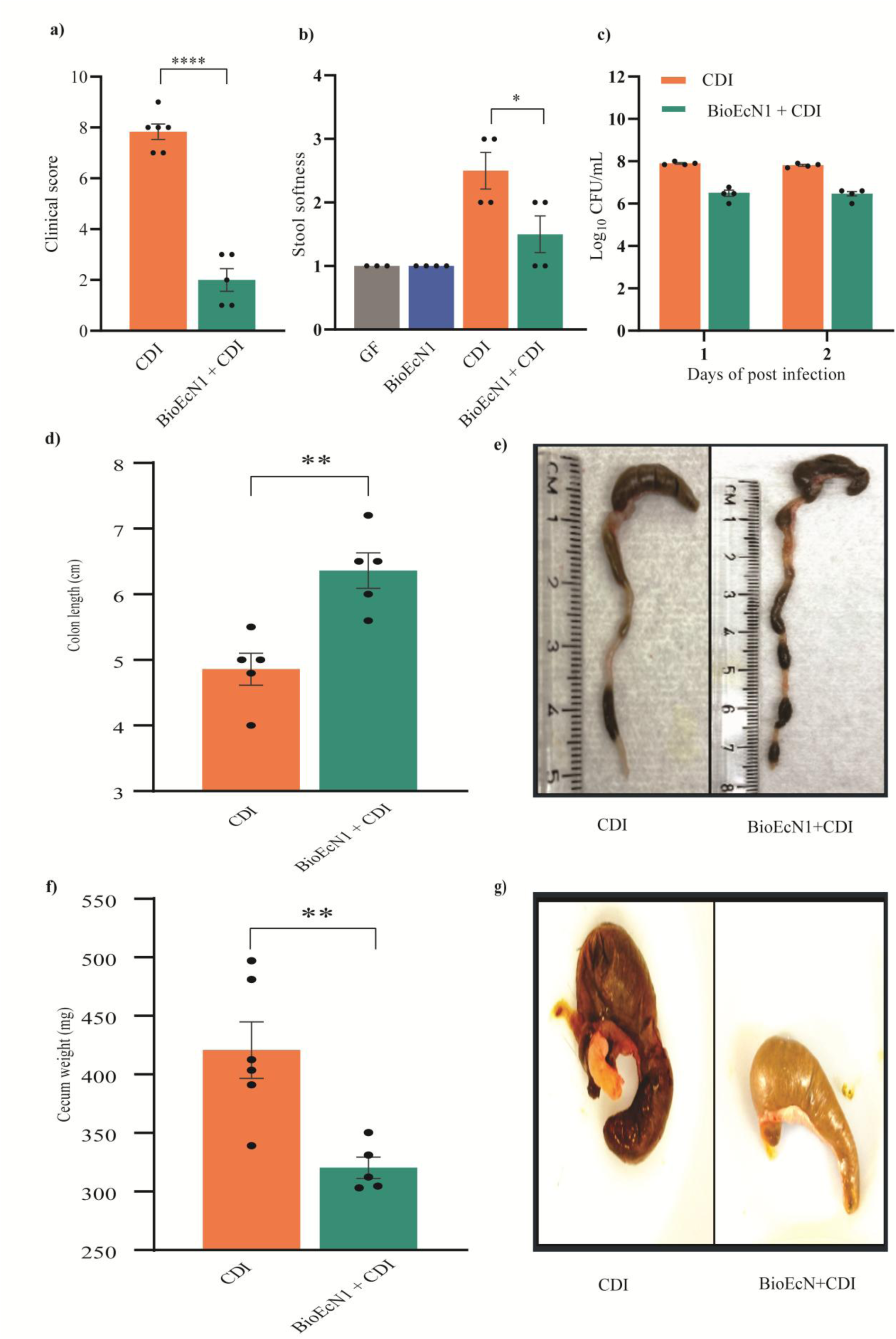
Clinical and intestinal outcomes following BioEcN1 prophylaxis in dysbiotic microbiota-associated mice. Mice harboring a human-derived dysbiotic microbiota received BioEcN1 before challenge with approximately 10⁷ CFU *C. difficile* strain 630. No antibiotic pretreatment was administered. (a) Clinical disease scores in CDI-only and BioEcN1-treated challenged mice. (b) Stool-consistency scores at [time] in uninfected controls, mice receiving BioEcN1 alone, CDI-only mice, and BioEcN1-treated challenged mice (c) Fecal *C. difficile* recovery on days 1 and 2 after challenge, expressed as log₁₀ CFU/mL of fecal suspension. (d) Colon length and (e) representative gross intestinal images at necropsy. Rulers indicate centimeters. (f) Cecum weight [specify whether contents were included] and (g) representative cecum images. Tissues were collected on day 7 after challenge.

## 4. Discussion

The main goal of this study was to determine whether prophylactic administration of CD27L-engineered EcN could attenuate CDI in mice. BioEcN1 enhanced inhibition of multiple *C. difficile* strains during coculture and reduced disease when administered before pathogen challenge. In germ-free mice, BioEcN1 reduced clinical disease, body-weight loss, and intestinal injury, with higher observed survival during the monitored period. Reduced disease was also observed in mice harboring a human-derived dysbiotic microbiota susceptible to *C. difficile* invasion. Together, these results support the feasibility of administering an engineered probiotic before pathogen exposure to reduce subsequent disease in susceptible mice.

The engineering strategy we used combines a genetically tractable probiotic chassis with a bacteriophage-derived antimicrobial payload. Previous work established the ability of recombinant CD27L to lyse *C. difficile*, providing a rationale for its production in a bacterial delivery system^35^. We adapted the plasmid-curing and pMUT-derived vector platform developed by Kan and colleagues, who demonstrated recombinant expression in the mouse intestine without antibiotic selection for plasmid maintenance^17^. In this study, we extend this platform to CD27L expression and prophylactic CDI protection. This approach complements other engineered-probiotic strategies, including the restoration of bile salt metabolism to limit *C. difficile* germination and growth^16^. Incorporating an endolysin offers a potential means of adding direct antibacterial activity to a probiotic chassis, although its activity and selectivity within the intestinal community require further characterization.

The *in vitro* findings establish enhanced pathogen inhibition as a functional outcome of EcN engineering. BioEcN1 reduced *C. difficile* recovery relative to parental EcN across all three tested strains, supporting its selection for prophylactic evaluation in mice. Administration before pathogen challenge reduced clinical disease and intestinal injury in germ-free mice, accompanied by lower fecal and cecal pathogen counts. The improvement in disease outcomes was more pronounced than the reduction in measured pathogen burden, demonstrating that protection occurred despite continued pathogen recovery. These findings support disease attenuation, rather than pathogen clearance, as the principal benefit of BioEcN1 prophylaxis under the tested conditions.

Maintaining an engineered construct without antibiotic selection is an important design requirement for a prophylactic probiotic. Administering antibiotics solely to retain a plasmid could disrupt the microbial community that the intervention is intended to protect. Kan et al. (2021) addressed this challenge by developing pMUT-derived vectors that were retained during passage through the mouse gastrointestinal tract without antibiotic selection^17^. In our study, stable BioEcN1 recovery on selective and nonselective media in monoassociated mice supported maintenance of the selectable construct over the observation period, extending the application of this platform to CD27L-based prophylaxis. Protection in mice harboring a human-derived dysbiotic microbiota further showed that BioEcN1 retained benefit in the presence of an established microbial community under the tested dosing conditions. Previous work has demonstrated that probiotic mucosal colonization varies among individuals and is associated with baseline host and microbiome features^36–39^. Testing BioEcN1 across additional donor-derived microbiota backgrounds would therefore help determine how interpersonal variation in community composition influences its recovery and prophylactic efficacy.

The protection observed in germfree and dysbiotic human microbiota associated mouse models provides a foundation for further development of BioEcN1 as a prophylactic intervention. Longer follow-up and systematic evaluation of dosing schedules could establish how long protection persists and whether it can be maintained after administration stops. Development of a marker-free construct, coupled with assessment of genetic stability, host responses, and effects on nontarget microorganisms, would advance its suitability for repeated use. The present findings also motivate testing in recurrence models^40,41^, where administration after standard CDI treatment could evaluate protection during a subsequent period of susceptibility^41,42^. Such studies would extend the current evidence for pre-challenge prophylaxis toward additional clinically relevant settings.

The broader implication our study is that an engineered probiotic could provide antimicrobial activity during a period of increased susceptibility, before clinical disease develops. Beyond CD27L, this approach offers a foundation for a tunable platform for intestinal delivery of protein-based antimicrobials such as endolysins. Future designs could adjust promoter control, expression, and release, while alternative payloads could expand the range of intestinal pathogens targeted. With further validation, such interventions could complement antibacterial treatment and microbiota-restoration strategies by providing protection during susceptible periods. The enhanced coculture inhibition and reduced disease observed in two mouse models provide preclinical support for this preventive strategy and a basis for developing probiotics with defined antimicrobial functions.

